# Regarding Emitter Positioning for Nanoflow Electrospray Ionization

**DOI:** 10.64898/2025.12.18.695161

**Authors:** Noah M. Lancaster, Scott T. Quarmby, Katherine A. Overmyer, Joshua J. Coon

**Affiliations:** Department of Chemistry, University of Wisconsin-Madison, Madison, WI, USA; Department of Biomolecular Chemistry, University of Wisconsin-Madison, Madison WI, USA; Morgridge Institute for Research, Madison, WI, USA

## Abstract

Nanoflow electrospray ionization is commonly used for proteomics due to its high sensitivity. Signal intensity, however, is dependent on optimal emitter positioning relative to the mass spectrometer inlet. Here, we characterize the effect of varied emitter positions on peptide signal intensity in all three dimensions using emitters and flows consistent with standard proteomic analyses. We observe improved signal robustness to x/y variations at increasing z distances and demonstrate that positioning within 1 to 2 mm of the optimal location will maintain consistent signal. Signal intensity behavior is consistent across the m/z range, suggesting a certain level of analyte-independence for proteomics analyses. These results provide insight for proteomics researchers using nanoflow LC-MS/MS.

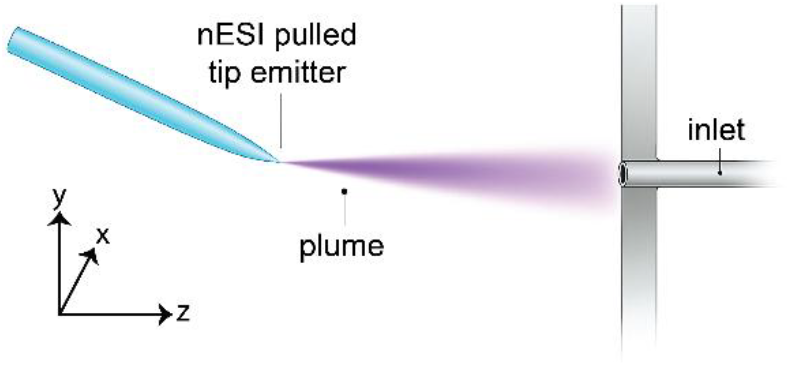

## INTRODUCTION

Owing to its high sensitivity, nanoflow electrospray liquid chromatography coupled with tandem mass spectrometry (nESI-LC-MS/MS) is widely used for protein sequence analysis.^1,2^ Whereas high-flow electrospray sources typically have relatively fixed emitter alignments and are more robust to changes in emitter positioning, nESI sources frequently require manual alignment in three dimensions.^3,4^ Since the number of ions reaching the MS system can vary depending on the location of the nESI emitter, positioning is a critical aspect of achieving robust and reproducible results.

Multiple reports illustrate the relationship between emitter position and electrospray signal intensity^2,4–10^; however, these studies typically examine a single dimension or evaluate emitters and flow regimes not typical of shotgun proteomic workflows. To understand these relationships, we investigate here the effect of emitter positioning on signal intensity when using fused-silica capillaries with integrated emitters at a flow rate of 300 nL/min, the standard setup across the field.^12^ Experiments were conducted using an Orbitrap Ascend mass spectrometer, which features a nanoflow source and asymmetric inlet capillary (high-capacity transfer tube, HCTT).^11^ This work represents, to our knowledge, the first report in the literature characterizing how emitter positioning impacts signal intensity for nanoflow electrospray ionization into an inlet capillary without radial symmetry.

Since most nESI emitters are manually positioned, understanding the precision in positioning that is required could assist in reducing batch-to-batch variation that is common in larger proteomic experiments. Further, recent efforts to improve the throughput of proteome analysis feature the use of multiple columns with multiple emitters aligned with the source at the same time.^13–17^ By performing measurements in all three dimensions, this report provides insight for implementing such a multi-emitter setup.

## EXPERIMENTAL SECTION

Emitter positioning measurements were performed by infusing BSA peptides at 300 nL/min into an Orbitrap Ascend mass spectrometer (Thermo Scientific) using a fused silica capillary (360 μm O.D, 75 μm I.D.) with integrated emitter (∼10 μm opening).^18^ The emitter was aligned to the inlet using the source camera and the x/y/z micrometer on a NanoSpray Flex source (Thermo Scientific). Note that x = 0 and y = 0 were defined as a point aligned to the center of the inlet capillary in these dimensions, and z = 0 corresponds to the plane orthogonal to the inlet entrance. Top-and side-view images of the starting position are shown in **Figure S1**. Raw Data and Code used for data analysis will be provided at the time of publication. Additional experimental and data analysis details are described in the **Supporting Information**.

## RESULTS AND DISCUSSION

We characterized signal intensity dependence on emitter position for nanoflow electrospray by infusing a mixture of peptides generated following trypsin digestion of purified BSA in an Orbitrap-quadrupole linear ion trap hybrid mass spectrometer (Orbitrap Ascend Tribrid). To ensure LC-MS applicability, we used an emitter size (∼ 10 micron orifice) and flow rate (300 nl/min) identical to those typical of capillary LC-MS.^18^ The resulting MS1 analysis produced numerous ions across the *m/z* range (**Figure S2**). We selected two of these ions, a doubly protonated species (LGEYGFQNALIVR at *m/z* 740.4) and a triply protonated species (RHPEYAVSVLLR at *m/z* 480.6), for their strong signal intensity and stability over the time range of the experiments here (**Figure S3** and **S4**). With the above setup, we tracked those m/z peaks as a function of emitter position in three dimensions and recorded intensity distributions (**Figure 1**). **Figure 1B** presents the ion signal across x-positions, where x = 0 is where the tip of the emitter is aligned with the center of the inlet and the dashed lines depict the MS inlet opening (0.6 mm, **Figure S5**).^19,20^ Strikingly, we observe that the signal is relatively stable and high across two millimeters of the x-dimension. Further, greater than 50% of the ion signal is retained at distances up to ∼five times the width of the inlet (i.e., full width half max (FWHM)). **Figure 1C** presents the same concept but in the y-dimension, where good signal stability is also observed across a similar length of two millimeters; however, the larger inlet opening in the y-dimension (1.6 mm) does not appear to proportionally impact the width of the intensity distribution. The ion plume is roughly symmetrical in both x- and y-dimensions^8^; therefore, it is expected that the smaller inlet width in the x-dimension would have a higher probability of being in the high density region of the plume than in the y-dimension. Proportional to inlet opening, the x position of the emitter would consequently have less of an effect on the intensity than the y position. We note the intensity distributions appear to be slightly off-center; however, upon fitting the data we find the centroids of the distributions (-0.11 and 0.26 mm, x- and y-respectively, **Figure S6**) were within the tolerance of the micrometer. **Figure 1D** presents the ion intensities as a function of emitter position in the z-dimension. Impressively, signal continues to be observed up to ∼7 mm removed from the inlet at levels close to 50% of the highest. Not surprisingly, the highest signal is observed at the closest position, but as noted above that signal gradually declines with increasing distance. Overall, identical trends were observed for both peptides over all dimensions.

**Figure 1.**
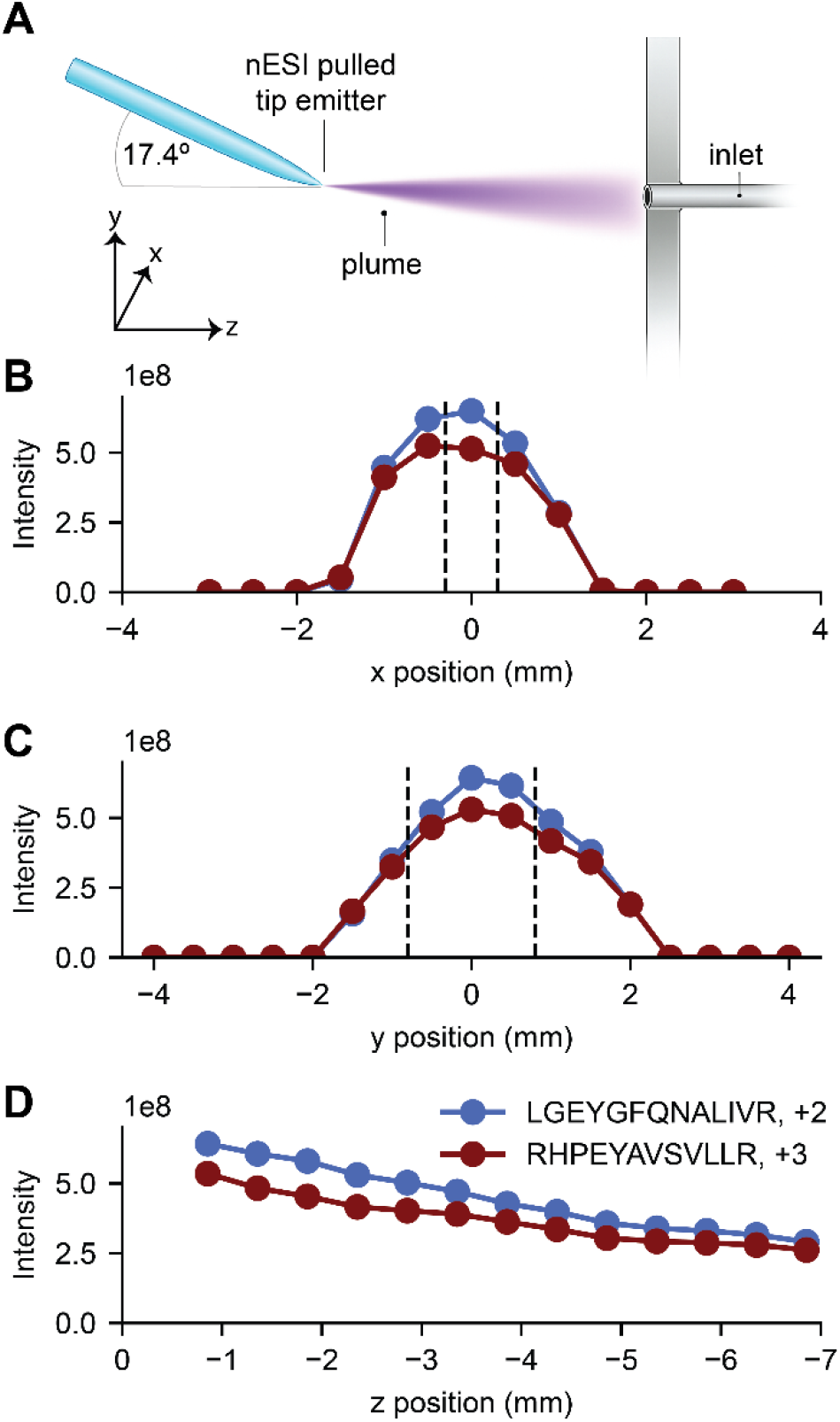
1D Positioning Experiments. (A) The x/y/z coordinate system used for this study. (B) Intensity dependence on x-position for two selected peptides (at y=0 and z = -0.856). Dotted lines indicate the estimated position of the inlet capillary edges. (C) Intensity dependence on y-position for two selected peptides (at x = 0 and z = -0.856). Dotted lines indicate the estimated position of the inlet capillary edges. (D) Intensity dependence on z-position for two selected peptides (at x = 0 and y = 0).

With these unidimensional observations complete, we next characterized the three-dimensional interplay of emitter position. For these experiments, we rastered the emitter across x, y, and z dimensions (**Figure 2** and **S7**). A key observation from these data is that signal intensity becomes more robust to x- and y-positioning as distance from the inlet increases (z-position). This observation is likely explained by a widening and flattening of the ion plume spatial distribution with increasing distance as previously reported.^9^ Specifically, with the emitter close to the inlet (z = -0.856 mm), the x, y tolerance to retain ∼90% signal is ∼1 mm, whereas at z = -4.856 mm the tolerance is ∼2-3mm (**Figure S8**). To our knowledge, these data are the first to map the three-dimensional space of the nESI plume in the context of the conditions that are typical in shotgun proteomics (i.e., capillary LC-MS). An especially exciting potential application of this knowledge is in the use of multiple emitters and/or parallel separations. In these cases, two or more emitters must be simultaneously positioned in front of the inlet, inhibiting the conventional alignment approaches where you put the emitter as centered and close as possible. Here, one desires to both achieve maximal ion signal while ensuring similar performance across the emitters. In such scenarios, our data suggest that moving 3-5 mm back in the z-dimension will ease the constraints on the x,y-positioning and reduce differences between the multiple emitters.

**Figure 2.**
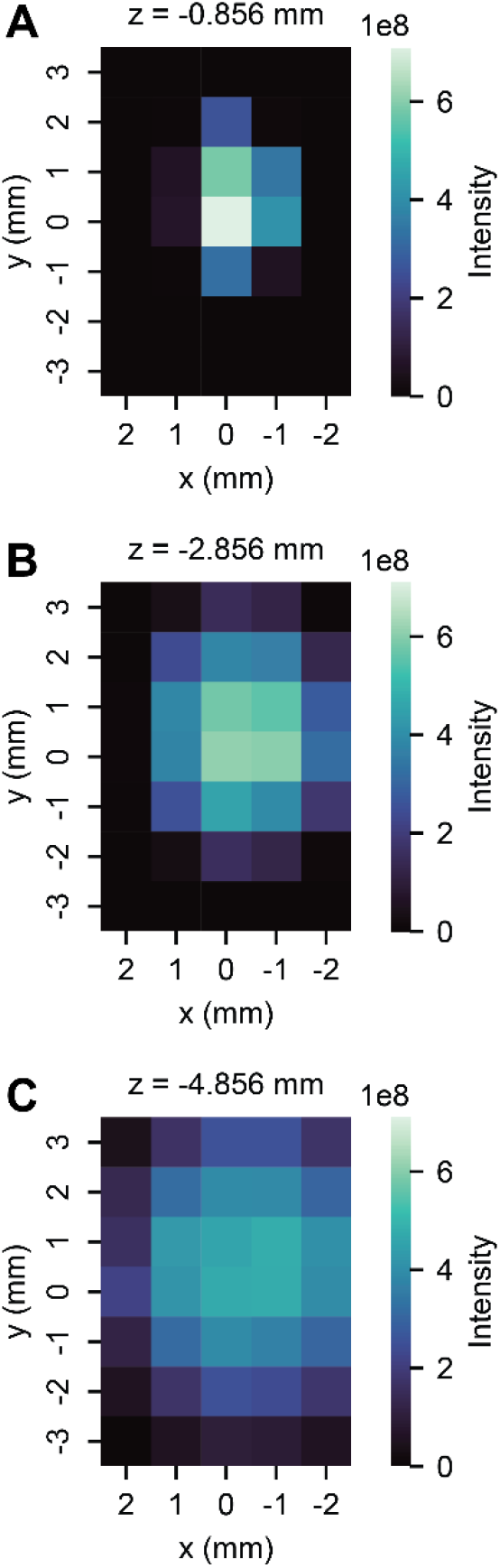
2D Positioning Experiments. Heatmaps showing intensity of LGEYGFQNALIVR (+2) at different x/y positions at (A) z = -0.856 mm, (B) z = -2.856 mm, and (C) z = -4.856 mm.

The source used in this study presented the emitter at an angle of ∼17.4° in the y-dimension (**Figure 1A** and **S1B**). From a purely geometric perspective, this angle could shift the y-dimension ion intensity distribution to increasingly positive values as the emitter is positioned further away in the z-dimension (see **Supplementary Note**). However, we find the centers of the y-dimension intensity distributions are consistent at different z-positions suggesting the angle had little to no impact (**Figure S9**). Gas flow into the inlet capillary has a collimating effect on the electrospray plume and is the dominating factor controlling ion trajectories.^21,22^ Additionally, the electrical potential difference between the emitter and the inlet capillary could curve the ion plume toward the inlet. These two effects likely explain why the geometric effect of the emitter angle is not observed.

Finally, to ensure there are no broader analyte specific effects at play, we looked at the effect of position across various *m/z* peaks (n = 243, charge > 1, *m/z* values ranging from ∼350 to ∼1250 *m/z*) stemming from the BSA tryptic digest. We plotted full-width half-maximum (FWHM) values for the x and y intensity distributions at the closest z position (**Figure 3A** and **3B**) and the intensity fold changes between the closest and furthest positions (*i*.*e*., z=-0.856 and z=-6.856, **Figure 3C**) vs *m/z*. Although there is variation in measured FWHM values and fold-change across analytes, this variation is not correlated with *m/z*. Note that outliers in these plots correspond to lower intensity peaks (**Figure S10**). We conclude that the effect of emitter positioning on signal intensity does not strongly depend on the analyte for peptide analyses.

**Figure 3.**
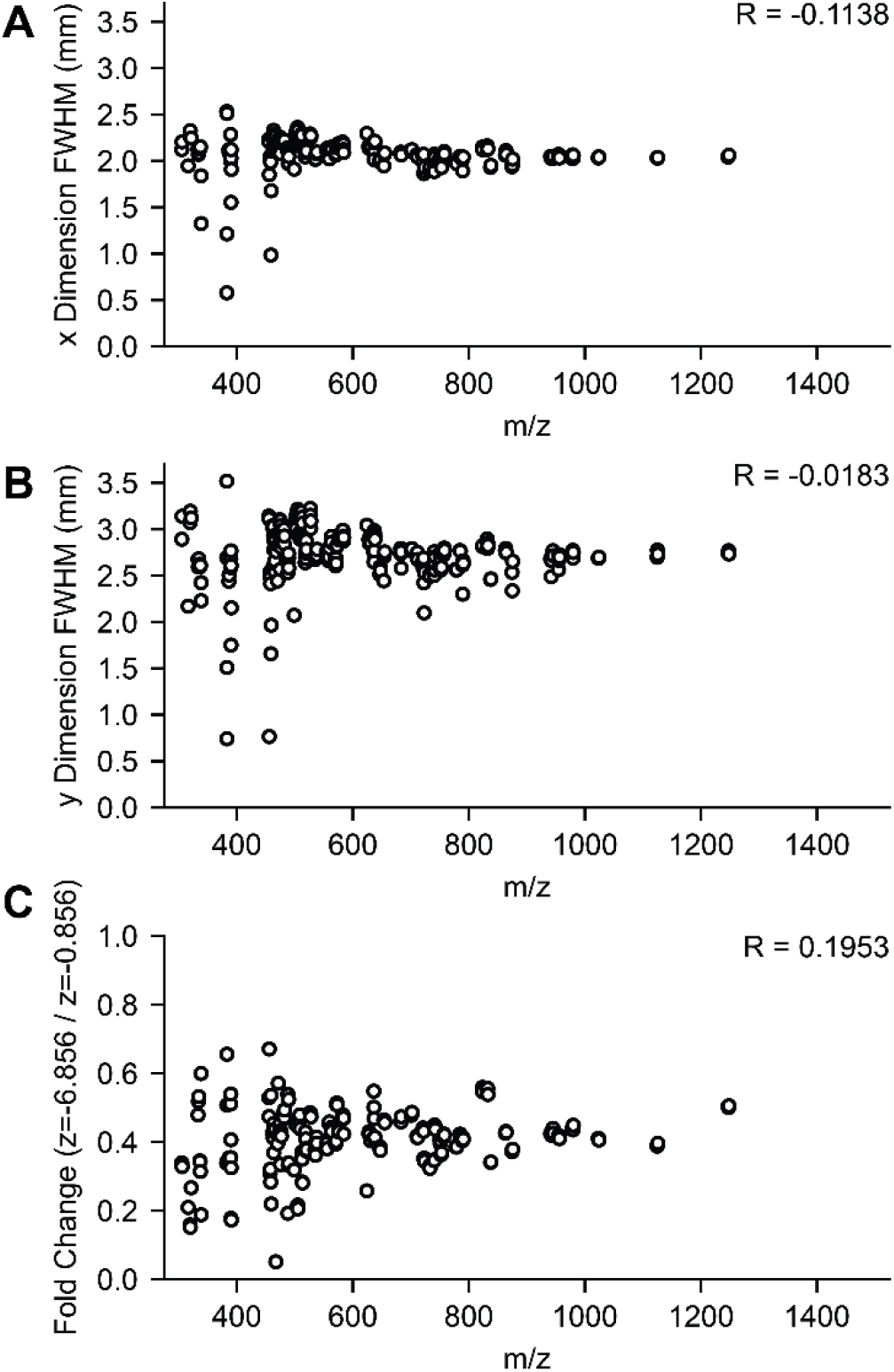
Dependence of Emitter Position Results on m/z. Mass spectra for the two extreme z-positions (at x = 0 and y = 0) were filtered for *m/z* peaks present in both with S/N>300 and charge >1. The full-width half-maximum (FWHM) was approximated via spline interpolation for the (A) x-distribution and (B) y-distribution at z = -0.856 mm, and (C) the fold change between the extreme z-positions was calculated (at x = 0, y = 0 mm). The Pearson correlation, R, of these values with m/z was calculated and is included at the top right of each panel.

## CONCLUSIONS

We report the dependence of peptide signal intensity on emitter positioning for nanoflow electrospray. Specifically, within 1-2 mm in any dimension, one can achieve reasonably consistent and robust signal. Distinct intensity profiles we observed here for the x- and y-dimensions likely arise from the asymmetric shape of the ion capillary. We demonstrate improved robustness of signal intensity to x/y variation at increasing z distances, an observation that will be helpful for positioning multiple emitters, for example.^13–17^

We provide evidence that the effect of the emitter position on signal intensity is analyte-independent for peptide analysis. This report provides insight into the role of emitter positioning on signal intensity for bottom-up proteomics and represents the first characterization of the effect of emitter position on electrospray signal intensity on an instrument with an inlet capillary lacking radial symmetry. Future areas of interest would be an examination of how trends vary across different flow rates or emitter types, as well as assessing how the exact intensity profiles vary across instruments with differences in their atmospheric interface.

## ASSOCIATED CONTENT

### Supporting Information

The Supporting Information is available free of charge on the ACS Publications website.

Supplementary results. Supplementary Methods, Supplementary Note, Figure S1 (Emitter Position Images Recorded with Source Camera), Figure S2 (MS1 Spectra of Infused BSA Tryptic Digest), Figure S3 (Signal Stability of Infused Peptides), Figure S4 (QC Measurements for Emitter Positioning Experiments), Figure S5 (Inlet Capillary Drawing), Figure S6 (Estimating Distribution Centroids), Figure S7 (2D Positioning Experiments for RHPEYAVSVLLR), Figure S8 (Width at 90% Maximum as Function of z Position), Figure S9 (Dependence of y Intensity Distribution on z Position), Figure S10 (Dependence of Emitter Position Results on Peak Intensity) (PDF)

## AUTHOR INFORMATION

### Corresponding Author

#### Author Contributions

NML prepared samples, collected data, and performed analysis. All authors contributed to writing and editing of the manuscript.

### Notes

J.J.C. is a consultant for Thermo Fisher Scientific, on the scientific advisory board for Seer, and a founder of CeleramAb Inc.

## ACKNOWLEDGMENTS

We are grateful for support from the National Institutes of Health (R35GM118110 to J.J.C.).

