## Supporting Information for "Regarding Emitter Positioning for Nanoflow Electrospray Ionization"

**Supplementary Methods**

*Materials and Reagents*

Ultrapure water was generated with a Barnstead GenPure Pro system (Thermo Scientific). Tris(2-carboxyethyl)phosphine hydrochloride (TCEP, C4706-2G), chloroacetamide (≥98%, C0267-100G), urea (U5378-1kg), and trifluoroacetic acid (TFA, HPLC grade, >99.9%) were obtained from Sigma-Aldrich. Tris Buffer (1 M Tris pH 8.0, 0.2 μm filtered, Invitrogen, AM9856), formic acid (LC-MS grade), acetonitrile (Optima LC/MS grade), and bovine serum albumin (BSA, Thermo Fisher Scientific, 23209, Albumin Standard) were obtained from Fisher Scientific. Trypsin (Sequencing Grade Modified Trypsin, V5113) was purchased from Promega. LysC (Lysyl Endopeptidase, 100369-826) was acquired from VWR.

*Sample Preparation*

BSA standard was diluted to 0.5 mg/mL with a final composition of 2 M Urea/100 mM Tris/2.5 mM TCEP/10 mM chloroacetamide. LysC was added at a 1:50 enzyme:BSA ratio and the sample was rocked at ambient conditions for 4 hours. Trypsin was added at a 1:50 enzyme:BSA ratio and the sample was rocked at ambient conditions overnight. Digestion was quenched with 10% TFA added to pH <2. The sample was centrifuged for 5 minutes at 9,000 g, desalted using a Strata-X 33 μm polymeric reversed phase SPE cartridge (Phenomenex), dried in a SpeedVac (Thermo Scientific), and resuspended in 0.2% formic acid. Peptide concentration was measured via NanoDrop (Thermo Scientific), and the sample was diluted to 0.5 mg/mL with a final solvent composition of 24% acetonitrile/0.2% formic acid.

*Data Collection*

Emitter positioning measurements were performed by infusing BSA peptides into an Orbitrap Ascend mass spectrometer (Thermo Scientific) with a Vanquish Neo UHPLC (Thermo Scientific). An emitter was prepared by laser pulling and etching a fused silica capillary (360 μm OD, 75 μm ID) with ~10 μm tip ID[1], connecting to the nanoLC, and aligned to the inlet with a NanoSpray Flex source (Thermo Scientific) and the source camera. The x = 0 and y = 0 points were defined as a point aligned to the center of the inlet capillary in these dimensions, and z = 0 was defined at the plane orthogonal to the inlet opening. **Figure S1** shows top and side-view images of the starting position**.** The emitter angle in the y-dimension was measured to be ~17.4° using the ImageJ software.[2] Infusion was performed by injecting 20-22 μL using Direct Control within Xcalibur at a flow rate of 300 nL/min and mobile phase conditions identical to the sample solvent composition. The infusion method provided over an hour of stable signal (**Figure S3**). For each set of positioning experiments, an infusion was begun, and the signal was monitored until it had stabilized (~15 min). Then, the emitter position was varied for data collection for no more than approximately 30 minutes to ensure stable spray. For this study, this procedure was performed once to collect data for the 1D experiments shown in **Figure 1**, and 3 times for the data in **Figure 2** (once at each z position). Measurements were taken at the starting positions throughout each data collection stage to confirm consistent signal intensity (**Figure S4**).

A spray voltage of 2000 V was applied with transfer tube temperature of 275°C. Orbitrap MS1 scans were acquired at 60,000 resolving power with scan range 300–1500 *m/z*, 30% RF Lens setting, 250% Normalized AGC Target, and maximum injection time of 100 ms. Signal intensity was allowed to stabilize after adjusting positions. For each position, 100 scans were acquired in the Tune software.

*Data Analysis*

An *in silico* BSA digest was performed using PeptideMass in Expasy[3] with two missed cleavages allowed. Outputs were used to select a doubly and triply charged precursor ($LGEYGFQNALIVR^{2+}$ and $RHPEYAVSVLLR^{3+}$) with high signal intensity and stability. Ion intensity was averaged across scans with a 15-ppm *m/z* tolerance using the 'pymsfilereader' library in Python. Calculation of approximate full-width half-maximum (FWHM) values for intensity distributions was performed by fitting the data with the 'UnivariateSpline' function within SciPy.[4] For the global comparisons across *m/z*, mass spectra for the two extreme z-positions (at x = 0 and y = 0) were averaged in Freestyle, followed by exporting peak lists and filtering peaks present in both spectra (within 5 ppm) for charge > 1 and S/N>300. This filtering retained higher intensity peaks more likely to be reliable indicators for the effect of emitter positioning on signal intensity. Pearson correlations were calculated using 'pearsonr' within SciPy.[4]

**Supplementary Note**

The Nanospray source used in this study exhibits a ~17.4° downward angle in the y dimension. An illustration of this geometry is shown below, where $\theta$ represents the downward angle, $d_z$ represents the distance of the emitter from the inlet in the z dimension, and $\Delta y$ represents the offset between the emitter's y position and the y position at the intersection of the central axis of the emitter and the plane of the inlet capillary.

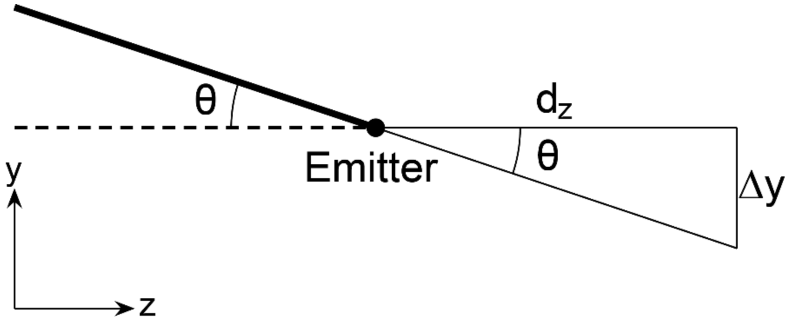


For an emitter angle, $\theta$, of 17.4° and an emitter distance, $d_z$, the offset, $\Delta y$, is defined by

$$d_z \tan\theta$$

Thus, for the z positions shown in **Figure 2** and **Figure S7**, the offsets are calculated as

| Absolute z Distance (mm) | Theoretical Δy (mm) |
|---|---|
| 0.856 | 0.268 |
| 2.856 | 0.895 |
| 4.856 | 1.522 |

This feature of the source geometry would be expected to impact the emitter positioning results in the y dimension if the geometric configuration of the source was a major driver of ion trajectories. However, this trend is not clearly observed as shown in **Figure S9**.

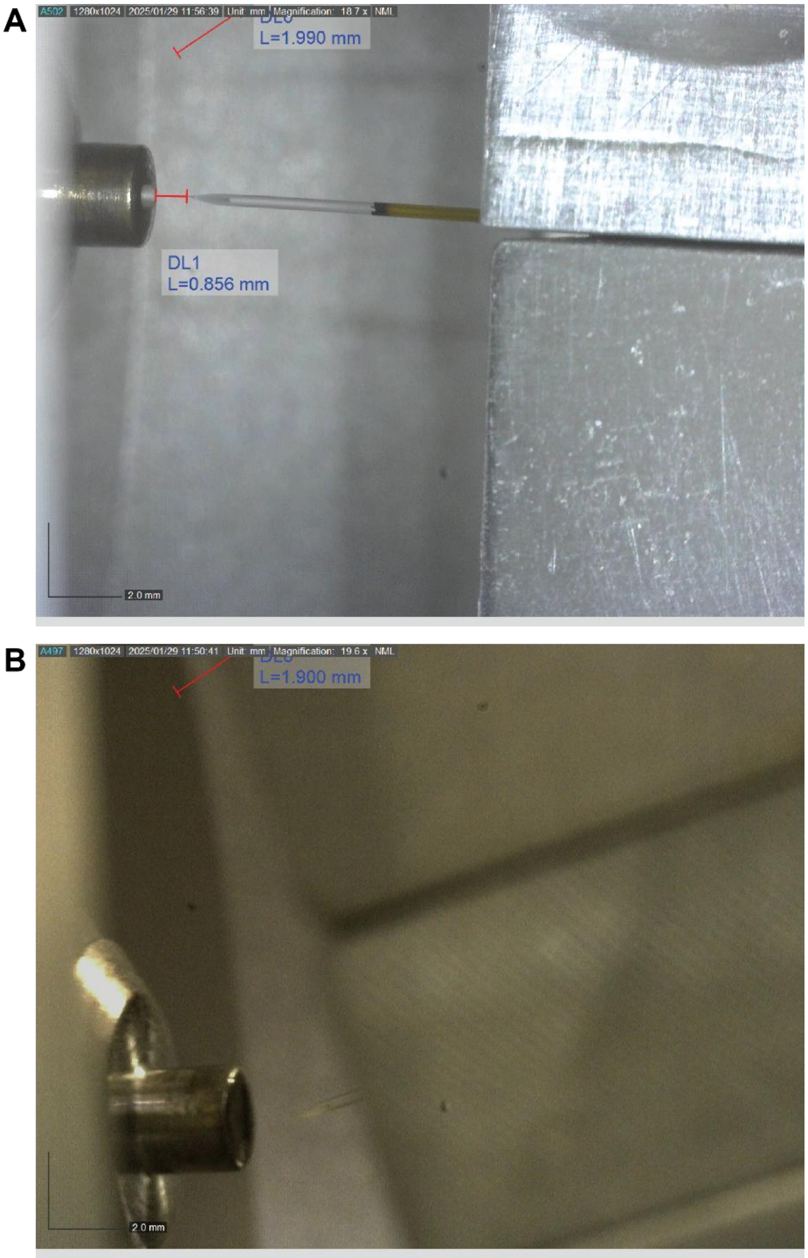


**Figure S1: Emitter Position Images Recorded with Source Camera.** (A) Top View of the emitter (x/z axes). The measured distance between the inlet capillary and the emitter tip was calibrated based on the known 360 µm O.D. of the silica capillary. Note that the 1.990 mm measurement is an artifact from the camera software. (B) Side view of the emitter (y/z axes). The angle of the emitter with respect to the inlet capillary was measured to be ~17.4° in ImageJ. Note that the 1.900 mm measurement is an artifact from the camera software.

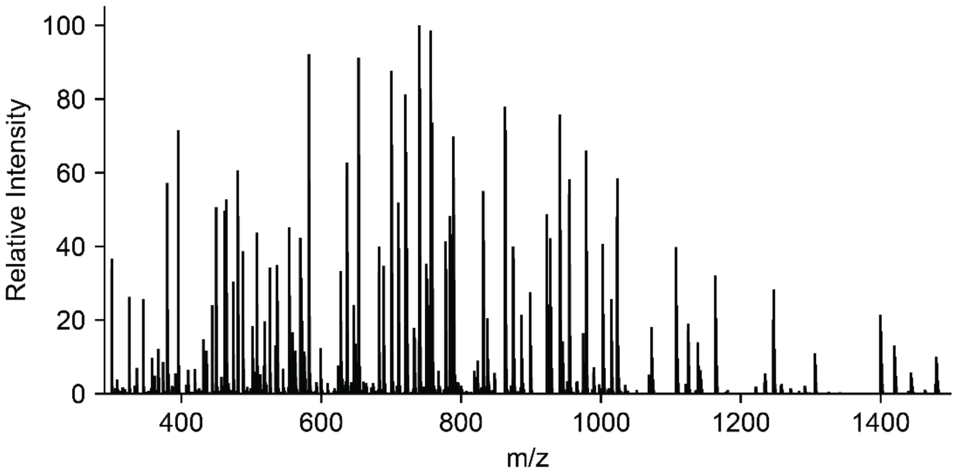


**Figure S2: MS1 Spectra of Infused BSA Tryptic Digest**. The Orbitrap MS1 spectra is shown for infusion of BSA peptides at x = 0 mm , y = 0 mm. z = -0.856 mm.

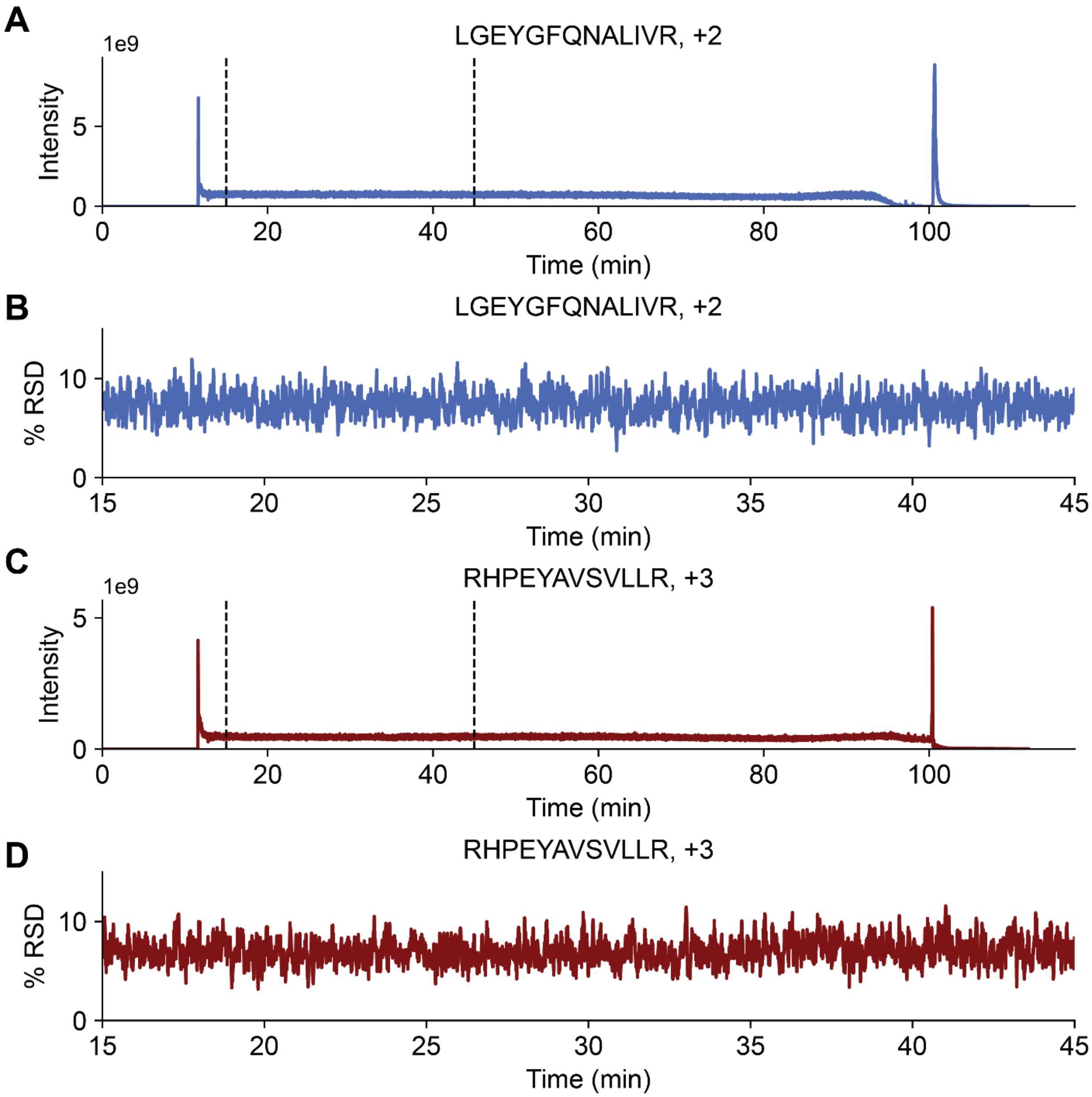


**Figure S3: Signal Stability of Infused Peptides.** (A) Signal stability over the infusion period for the LGEYGFQNALIVR$^{2+}$ ion. Dashed lines indicate the typical time window when emitter positioning data was collected during the sample infusion. (B) Plot showing the rolling %RSD value for the LGEYGFQNALIVR$^{2+}$ signal within the typical data collection window indicated in (A). (C) Signal stability over the infusion period for the RHPEYAVSVLLR$^{3+}$ ion. Dashed lines indicate the typical time window when emitter positioning data was collected during the sample infusion. (D) Plot showing the rolling %RSD value for the RHPEYAVSVLLR$^{3+\ +}$ signal within the typical data collection window indicated in (C). Rolling %RSD values were calculated over 20 scans.

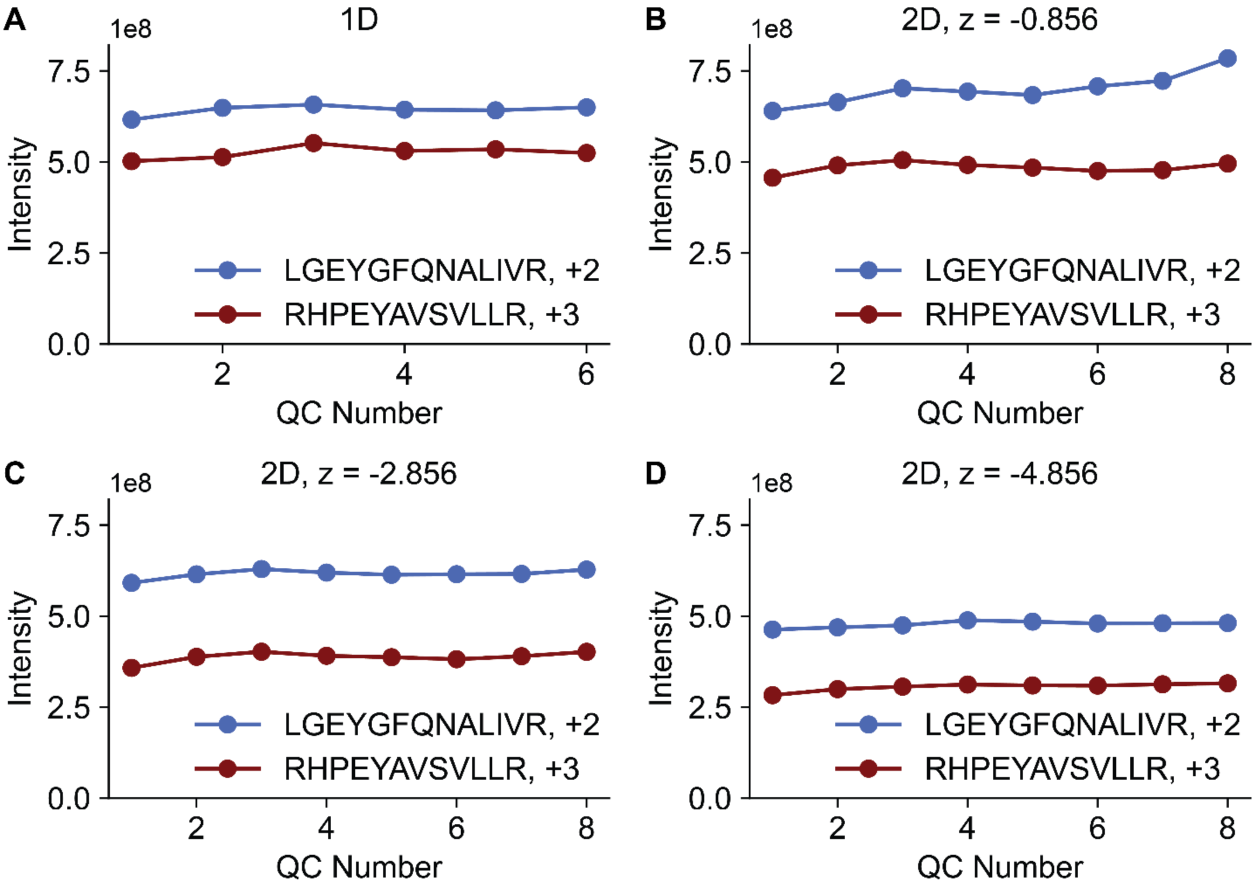


**Figure S4. QC Measurements for Emitter Positioning Experiments.** The position was returned to the starting position for each experiment periodically throughout data collection to confirm signal stability during the experiments. The signal intensity is shown for the (A) one-dimensional experiments shown in **Figure 1**, and the two-dimensional experimental shown in **Figure 2** and **Figure S7** at (B) z = -0.856, (C) z = -2.856, and (D) z = -4.856 mm. For all measurements, x = 0 mm and y = 0 mm.

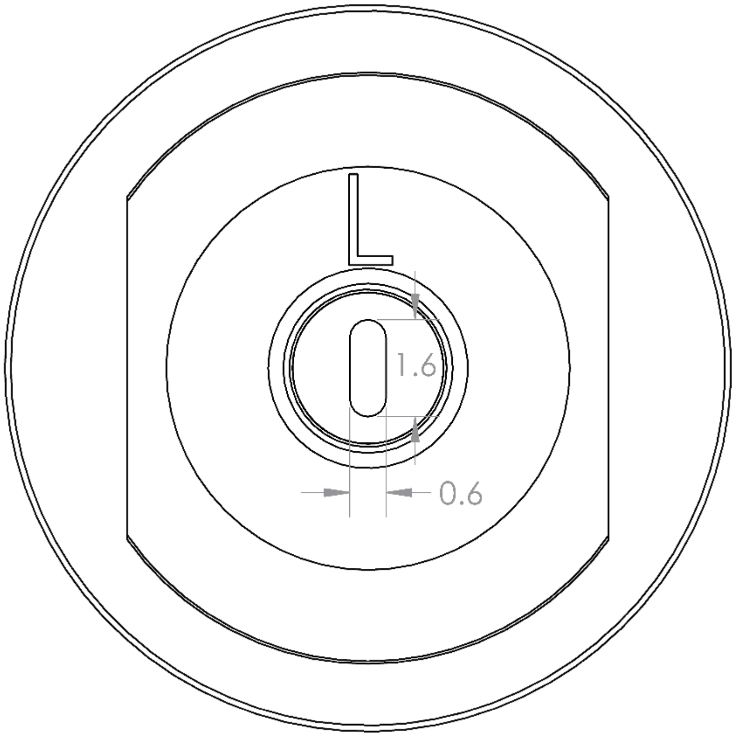


**Figure S5. Inlet Capillary Drawing.** A drawing of the high-capacity transfer tube (HCTT) is shown. Measurements are reported in mm with 0.6 mm being the width in the x dimension and 1.6 mm being the width in the y dimension.

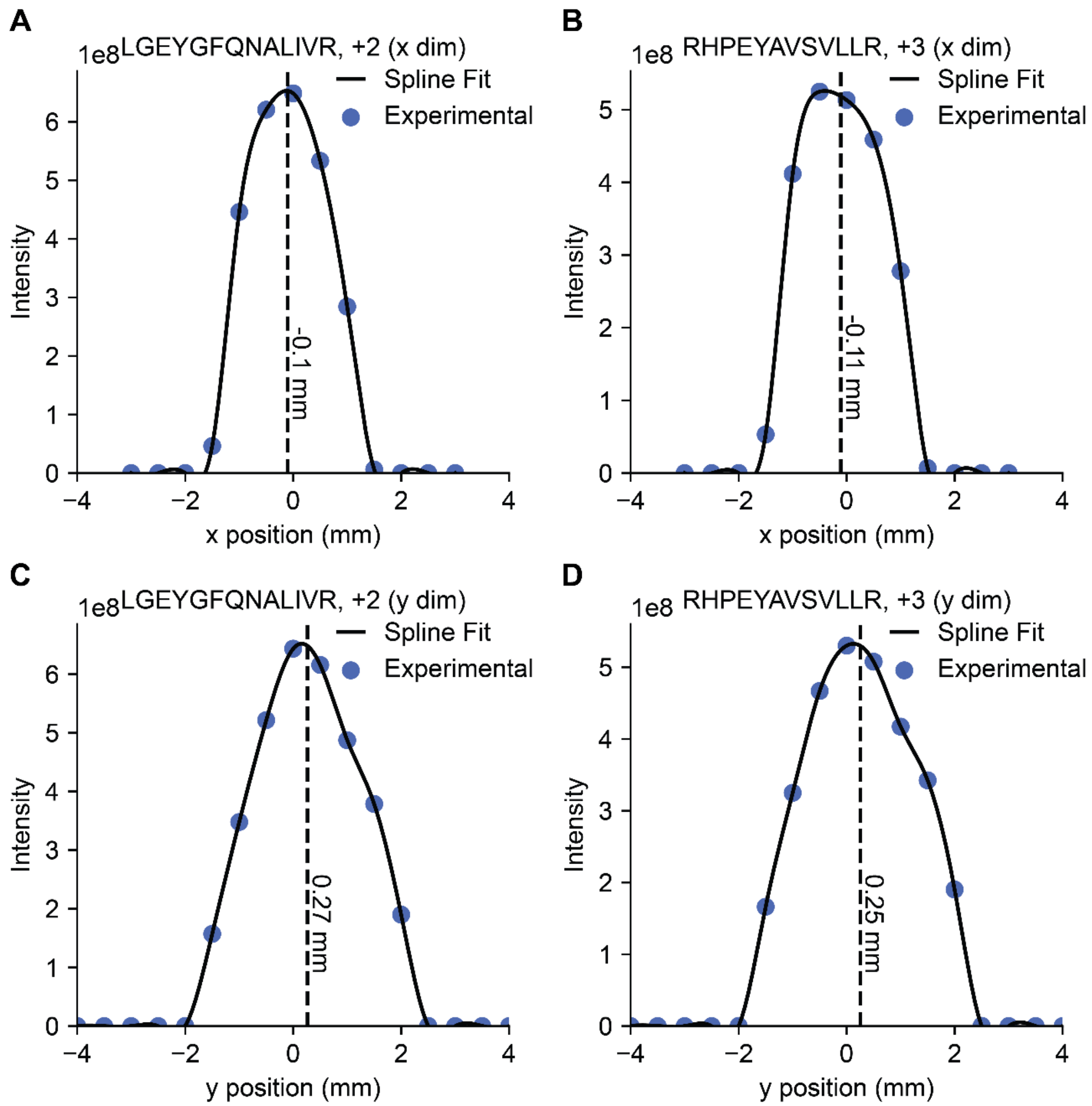


**Figure S6**. **Estimating Distribution Centroids.** For the data shown in **Figure 1B** and **1C**, the centroids of the distributions in the x- and y-dimensions (at z = -0.856) were approximated by fitting the data to a spline interpolation function with SciPy. Data are shown for (A) LGEYGFQNALIVR (+2) for the x dimension, (B) RHPEYAVSVLLR (+3) for the x dimension, (C) LGEYGFQNALIVR (+2) for the y dimension, and (D) RHPEYAVSVLLR (+3) for the y dimension. Dotted black lines with text annotations indicate the position of the centroid for the spline function calculated as $\bar{x} = \sum(x_i\ I_i)/\sum I_i$, where $x_i$ is an x position in the spline fit and $I_i$ is the corresponding intensity value at that position.

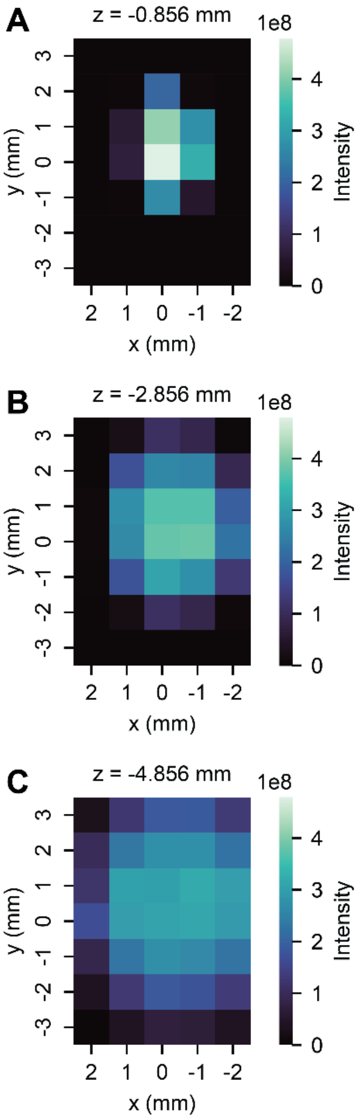


**Figure S7. 2D Positioning Experiments for RHPEYAVSVLLR.** Heatmaps showing the intensity of RHPEYAVSVLLR (+3) at different x/y positions at (A) z = -0.856 mm, (B) z = -2.856 mm, and (C) z = -4.856 mm.

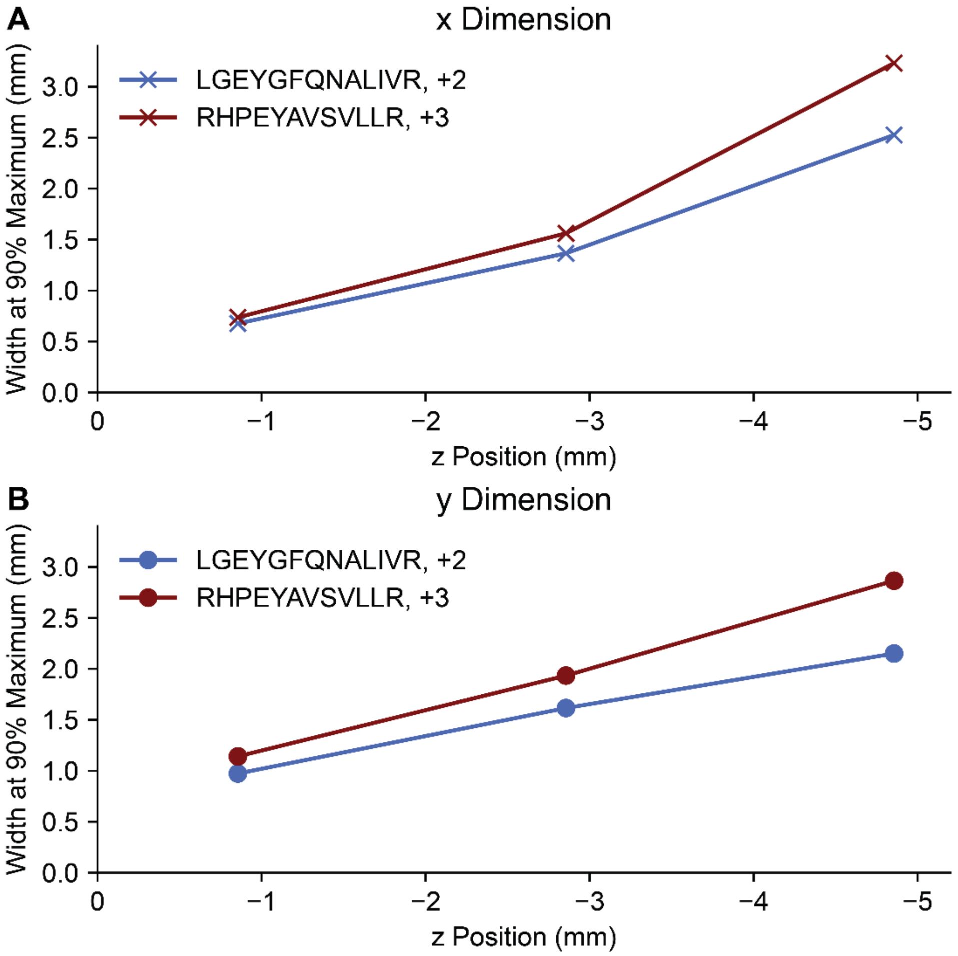


**Figure S8**. **Width at 90% Maximum as Function of z Position.** For the data shown in **Figure 2** and **S7**, the width at 90% maximum of the intensity distribution in the x- and y-dimensions were approximated by fitting the data to a spline interpolation function with SciPy. The approximate width values for the x- and y-distributions are then shown for the two different precursors ions as a function of the z position.

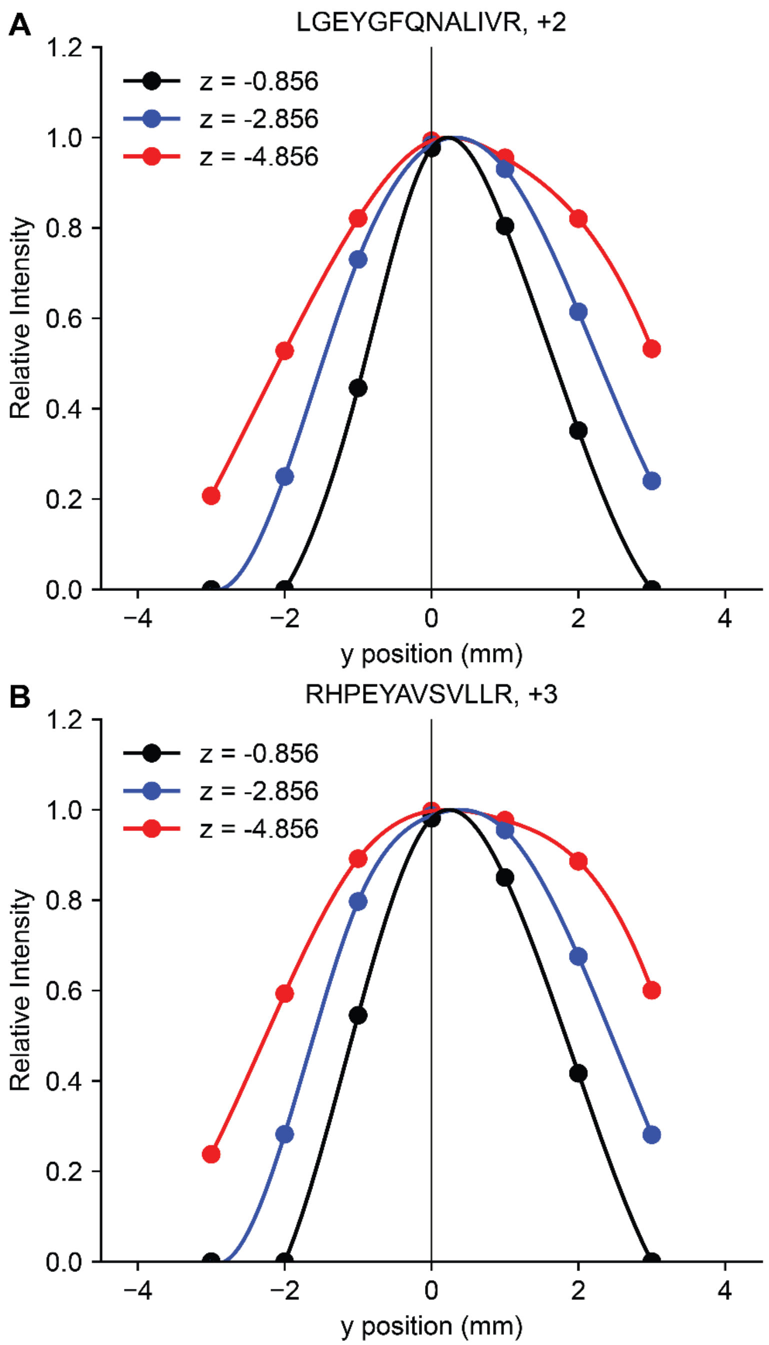


**Figure S9. Dependence of y Intensity Distribution on z Position.** For the data shown in **Figure 2** and **S6**, the y position-dependent normalized intensity distributions (at x = 0) were fit to a spline interpolation function with SciPy. Data are shown at different z positions for (A) LGEYGFQNALIVR (+2) and (B) RHPEYAVSVLLR (+3). The y = 0 point is indicated with a black line. Dots represent the experimental datapoints and the lines represent the spline function.

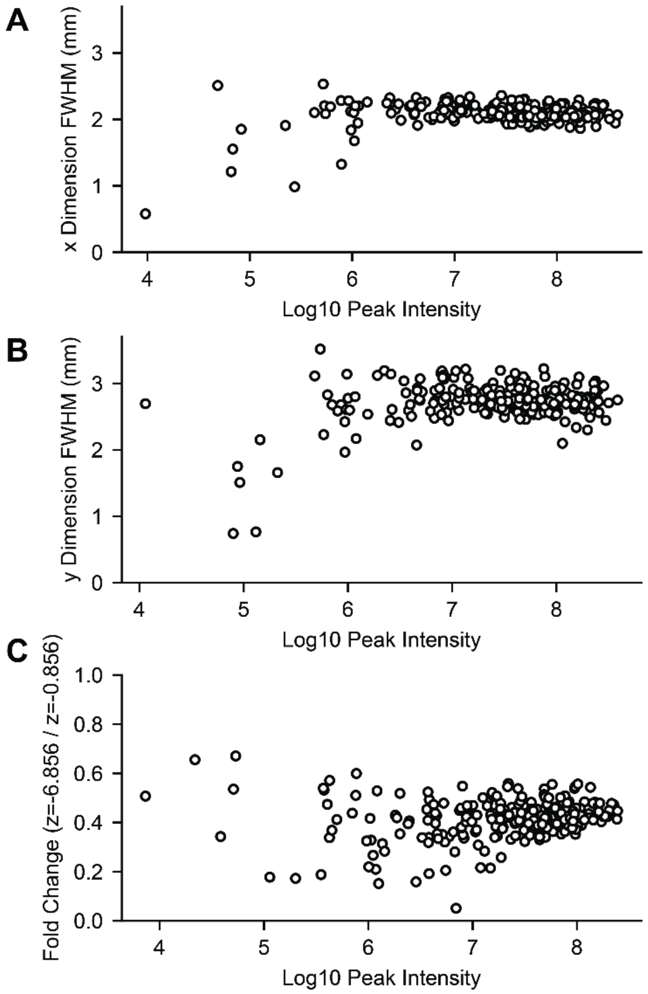


**Figure S10. Dependence of Emitter Position Results on Peak Intensity** Mass spectra for the two extreme z-positions (at x = 0 and y = 0) were filtered for m/z peaks present in both spectra with S/N > 300 and charge > 1. The full-width half-maximum (FWHM) was approximated via spline interpolation for the (A) x-distribution (at z = -0.856 mm) and (B) y-distribution (at z = -0.856 mm), and (C) the fold change between the extreme z-positions (at x = 0, y = 0 mm) was calculated. These data show that low intensity peaks exhibit much greater variation.
